# Modulation of Karyopherin Levels Attenuates Mutant Ataxin-1-Induced Neurodegeneration

**DOI:** 10.1101/2023.07.12.548780

**Authors:** Elena K Ruff, Dylan Lawrence Timperman, Adulfo Anaya Amador, Isabella Aguirre Lamus, Khondker Salim, Maria de Haro, Ismael Al-Ramahi

## Abstract

Neurodegenerative diseases are characterized by the abnormal accumulation of disease-driving proteins. Emerging evidence suggests that nucleocytoplasmic transport (NCT) components play a critical role in neurodegeneration. This study investigates the effects of modulating the levels of different karyopherins on mutant Ataxin-1 (mATXN1)-induced neurodegeneration in Spinocerebellar Ataxia Type 1 (SCA1) *Drosophila* and cell models. Our findings reveal that ATXN1 [82Q] interacts with KPNAs in the nucleus and cytoplasm of neurons. Increasing KPNA levels ameliorates ATXN1 [82Q]-induced neurodegeneration and progressive neuronal dysfunction. Surprisingly, elevated KPNA levels did not increase nuclear mATXN1, instead, mechanistic analyses demonstrate that KPNA retains mATXN1 in the cytoplasm, reducing its nuclear accumulation. Moreover, higher KPNA leads to a decrease in soluble oligomeric mATXN1. Interestingly, inhibition of a different karyopherin, KPNB1, elevated KPNA levels and reduced nuclear mATXN1 in human neuronal precursor cells. Consistently, knockdown of KPNB1 attenuates ATXN1 [82Q]-induced neurodegeneration and reduces its nuclear aggregation in *Drosophila*. These results indicate that KPNAs may act as chaperones for mutant ATXN1, preventing its nuclear translocation and reducing its pathological effects. Importantly, they also constitute a proof of principle that retaining mATXN1 in the cytoplasm represents an attractive and viable therapeutic option. Given the dysregulation of karyopherins in many neurodegenerative diseases and their emerging role as chaperones, the results presented here may extend beyond SCA1 into other disorders like Alzheimer’s or Parkinson’s disease.

## Introduction

A common hallmark of neurodegenerative diseases, though characterized by a number of different cognitive, psychological, or motor clinical manifestations, is the abnormal accumulation of disease-driving proteins ^1,2^. The majority of neurodegenerative diseases progress in an irreversible manner and currently have no effective treatments ^3^.

Growing evidence indicates that nucleocytoplasmic transport (NCT) plays a key role in neurodegenerative diseases ^4^. For instance, impaired NCT is identified as a major disease mechanism in both Amyotrophic Lateral Sclerosis (ALS) and Frontotemporal lobar degeneration (FTLD). TAR DNA-binding protein 43 (TDP-43), the disease protein identified in 97% of ALS and FTLD cases, mislocalizes and aggregates in the cytoplasm of neurons ^5–9^. These TDP-43 aggregates also contain Nucleoporins (Nups) and NCT factors, contributing to further dysregulation of NCT ^10^. Evidence suggests that NCT dysregulation is a major mechanism in AD as well ^4,11^. Invaginations and abnormalities of the nuclear envelope, in conjunction with abnormal localization of importins, Nups and RanGDP in AD patient brains add to this growing evidence ^4,10–16^ Karyopherins (KPNs), i.e., importins, exportins, and transportins, are the family of proteins that facilitate nuclear transport. Alpha KPNs (KPNAs) bind nuclear localization signal (NLS) sequences and then, along with their cargo, are carried by Beta KPNs (KPNBs) into the nucleus. This process also necessitates Nups, Ran, and RanGTPases ^17–19^. There are several human KPNs, each of which associates with a specific set of cargos^4^. Many of these KPNs are dysregulated in ALS, FTD, AD, and Huntington’s Disease (HD) ^12,20–24^. Interestingly, importins, also called nuclear import receptors (NIRs), can reduce aggregation of TDP-43 and other RNA binding proteins in cell and animal models by acting as chaperones for the toxic, aggregated protein ^25,26^. It has been shown that KPNs can disaggregate TDP-43, increase its nuclear localization and mitigate its toxicity ^25–27^.

Spinocerebellar Ataxia Type 1 (SCA1) is one of 9 Polyglutamine (polyQ) diseases caused by the expansion of a translated CAG tract in the *ATXN1* gene ^28^. This polyQ expansion causes a toxic gain of function in the protein, which misfolds and accumulates in the nuclei of neurons ^29^. Notably, polyQ expansions can be understood as a type of low complexity domain (LCD) or Prion Like Domain (PrLD) ^30^. ATXN1 itself has also been identified as a PrLD ^27^.

ATXN1 aggregates induce their pathogenic effect when they localize to the nucleus where they form nuclear bodies (NBs) ^30,31^. The removal of polyQ-expanded ATXN1’s functional NLS prevents disease phenotypes in SCA1mouse models. ^31,32^. This indicates that nuclear localization of mATXN1 is required for pathogenesis. Interestingly, the ATXN1 protein interactome includes many nuclear transport components ^33^. Additionally, the presence of polyQ-ATXN1 disrupts the normal NCT mechanisms. This includes inhibition of importins and exportins from binding their normal cargo, reducing the availability of RanGDP, and disrupting the nuclear pore complex ^33^.

Here we explore the functional consequences of modulating karyopherin levels on mATXN1-induced neurodegeneration. We show that ATXN1 [82Q] interacts with both *Drosophila* and human KPNAs in the nucleus and in the cytoplasm of neurons. Moreover, we show that elevated KPNA levels mitigate ATXN1 [82Q]-induced neurodegeneration and progressive neuronal dysfunction. Surprisingly, mechanistic analyses reveal that increased KPNA expression promotes the cytoplasmic retention of ATXN1 [82Q] and reduces its nuclear accumulation in the *Drosophila* central nervous system (CNS). Additionally, we observe that an increase in KPNA levels leads to a decrease in soluble oligomeric ATXN1 levels. Interestingly, pharmacological inhibition of KPNB1 increases cytoplasmic levels of KPNAs and decreases nuclear levels of mATXN1. Furthermore, knockdown of KPNB1 ameliorates ATXN1 [82Q]-induced neurodegeneration and lowers mATXN1 nuclear aggregates. Together, our findings support a growing body of evidence suggesting a molecular chaperone-like role for NIRs and provide convincing evidence that retention of mATXN1 in the cytoplasm by modulation of karyopherin levels holds therapeutic potential for SCA1 treatment. Given the pervasive role KPN dysregulation plays in many neurodegenerative disorders ^20^, the results presented here may extend beyond SCA1 and elucidate common mechanisms across multiple neurodegenerative diseases, such as AD or Parkinson’s disease (PD).

## Results

### KPNAs colocalize with ATXN1 [82Q] aggregates in the nucleus and cytoplasm of neurons

The ATXN1 interactome includes several NCT proteins, including KPNB3 and KPNA2 ^33^, but the functional consequence of these interactions has not been fully characterized. In order to investigate the potential interaction between ATXN1 [82Q] and KPNAs *in vivo*, we expressed ATXN1 [82Q] alongside HA-tagged nuclear import receptors using the *nrv2*-*GAL4* central nervous system (CNS) driver in *Drosophila* ventral nerve cords (VNCs). Immunofluorescence was performed in order to visualize the protein localization. In control CNS, the ATXN1 [82Q] signal is found predominantly in the nucleus (Fig. **1B**). Co-expression of ATXN1 [82Q] alongside Kap-α1 (the *Drosophila* homolog of KPNA6) results in the two proteins colocalizing (Fig. **1C**). Interestingly, the signals from ATXN1 [82Q] and the *Drosophila* KPNA overlap in distinct aggregate shapes in both the nucleus and the cytoplasm (Fig. **1C**). A similar colocalization in the nucleus and cytoplasm is also observed when ATXN1 [82Q] is co-expressed with human KPNA5 (Fig. **1D**) or KPNA2 (Fig. **1E**) in the *Drosophila* CNS. These results indicate that ATXN1 [82Q] interacts with *Drosophila* and human KPNAs *in vivo* and that this interaction happens both in the nucleus and the cytoplasm of neurons.

**FIG. 1.**
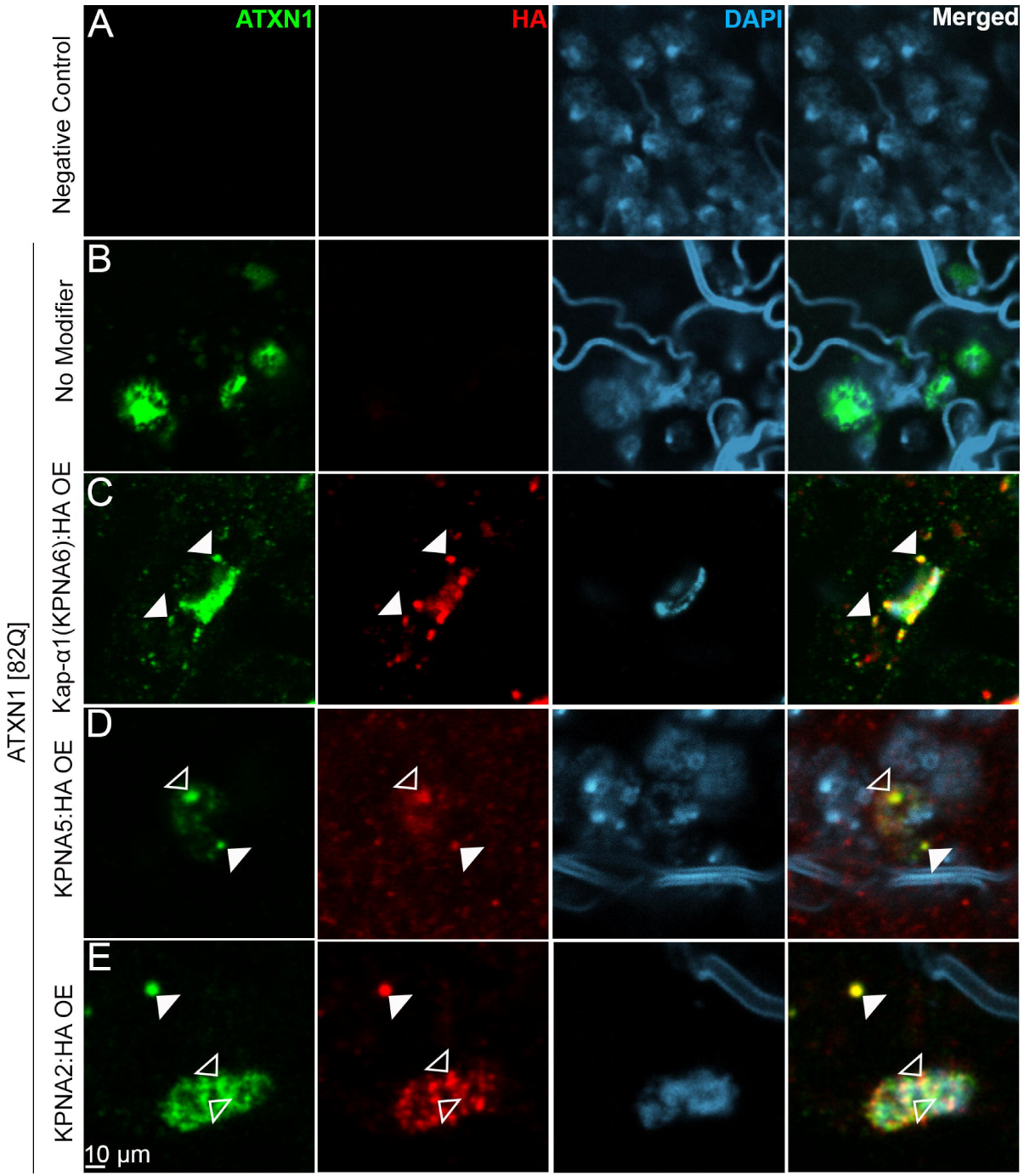
KPNAs interact with ATXN1 [82Q] in the nucleus & the cytoplasm in the *Drosophila* CNS. Confocal images of *Drosophila* neurons in the adult ventral nerve cord (VNC) stained with anti-ATXN1 antibody (green), antibodies staining different KPNA proteins (red) and DAPI marking the nuclei (blue). (**A**) No ATXN1 [82Q] (green) or KPNA staining (red) is detected in negative controls. (**B**) ATXN1 [82Q] staining (green) detected in discrete aggregates. (**C - E**) Reveal colocalization of ATXN1 [82Q] with either *Drosophila* Kap-α1 (**C**) or human KPNA5 (**D**) and KPNA2 (**E**). Notice the overlap of the ATXN1 [82Q] and KPNA signals in both the nucleus (*hollow arrows*) and the cytoplasm (*solid arrows*) in the merged images. A-E scale bars = 10μm. Flies were raised at 28°C. Transgenes were expressed from the *nrv2-GAL4* driver.

### Increased KPNA levels ameliorate ATXN1 [82Q]-induced neurodegeneration in Drosophila

Following our findings that KPNAs colocalize with mATXN1, we investigated the functional significance of this interaction. First, we used the *gmr*-*GAL4* driver for retinal ATXN1 [82Q] expression, which leads to external (Fig. **2B**) and internal (Fig. **2B’**) degenerative phenotypes. Interestingly, co-expression of ATXN1 [82Q] together with the *Drosophila* homologs of either KPNA6, KPNA4, or KPNA2 ameliorates the ATXN1 [82Q]-induced neurodegeneration (Fig. **2C-F; 2C’-F’**). The attenuation of the ATXN1 [82Q] phenotypes was robust both when analyzing external eye morphology (Fig. **2C-F**) as well as when investigating the internal photoreceptor structure (Fig. **2C’-F’**). Co-expression of ATXN1 [82Q] together with human KPNA2 and KPNA5 also ameliorated ATXN1 [82Q]-induced neurodegeneration (Fig. **2G-H; 2G’-H’**). To corroborate the genetic interaction between ATXN1 [82Q] and KPNAs in a time progressive assay, we analyzed the effect of co-expressing KPNAs together with ATXN1 [82Q] in the CNS using a behavioral test. Expression of ATXN1 [82Q] in the *Drosophila* CNS leads to progressive dysfunction which can be quantified using the motor performance (speed) of the animals as a function of age using a climbing assay (Fig. **2I-L**) (Gray lines). We have used this assay to analyze toxic protein effects on neurons in other *Drosophila* models of neurodegenerative diseases ^34,35^. Expression of ATXN1 [82Q] in the *Drosophila* CNS results in a progressive decline in motor function over time, compared to negative controls (Fig. **2I-L**) (Blue lines). Increasing the levels of either *Drosophila* or human KPNAs in the central nervous system significantly ameliorates this progressive ATXN1 [82Q]-induced neuronal dysfunction (Fig. **2I-L**) (Red lines).

**FIG. 2.**
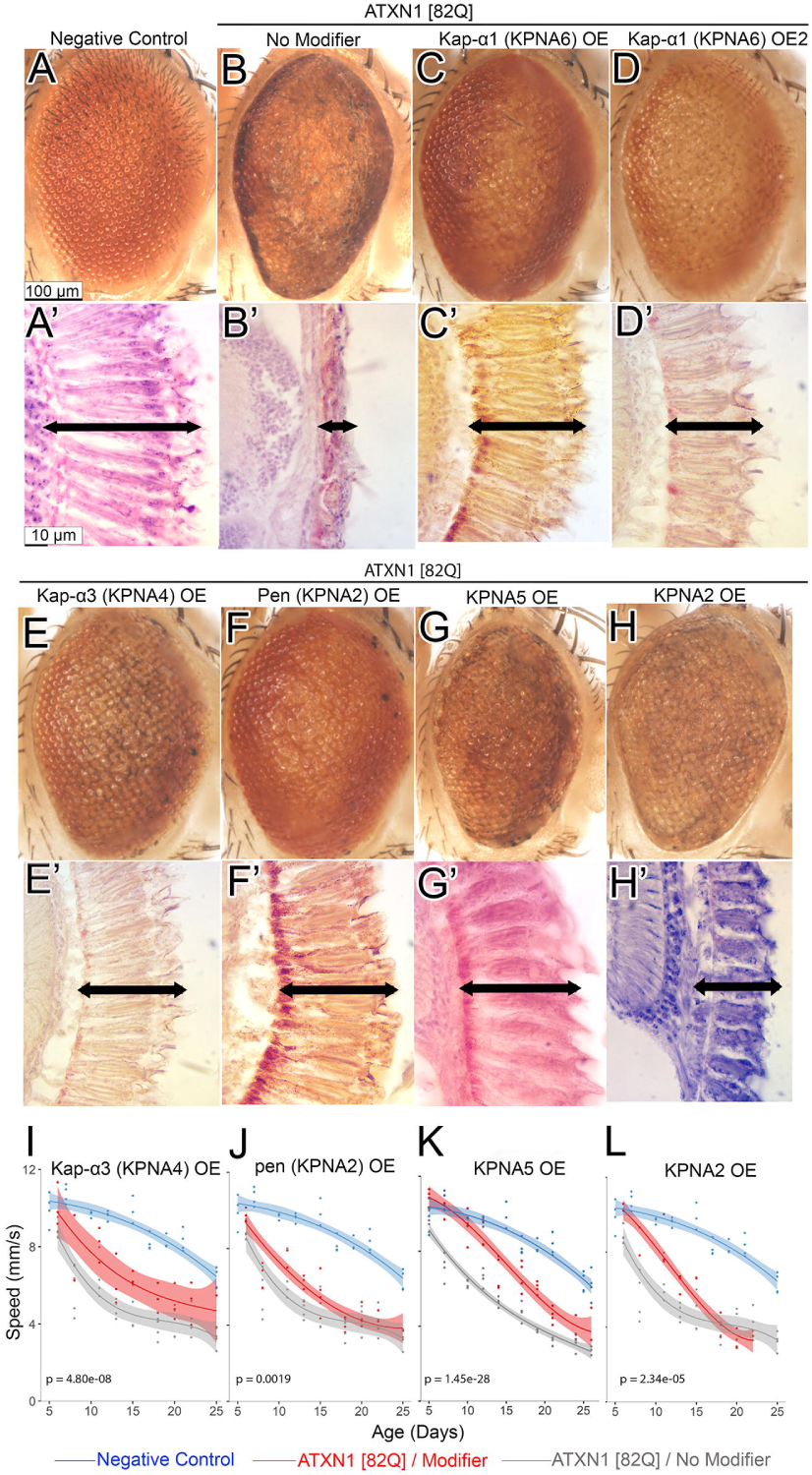
Increased levels of KPNAs ameliorate ATXN1 [82Q]-induced photoreceptor degeneration and neuronal dysfunction. (**A-H**) External eye images and **(A’-H’**) retinal paraffin sections of the indicated genotypes. Arrows indicate photoreceptor length. (**A, A’**) Control eyes display organized ommatidia, evenly spaced interommatidial bristles, and long, straight, parallel photoreceptors. (**B, B’**) ATXN1 [82Q] eyes exhibit disorganized ommatidia, uneven or missing bristles, and fewer, shorter, and curved photoreceptors. (**C-H; C’-H’**) Increased *Drosophila* (**C-F, C’-F’**) or human (**G-H, G’-H’**) KPNA levels improve ATXN1 [82Q]-induced external and retinal degeneration phenotypes. ATXN1 [82Q] / KPNA eyes show better ommatidial arrangement (**C-H** vs. **B**) and longer, straighter photoreceptors (**C’-H’** vs. **B’**). Scale bars: A-H = 100 μm, A’-H’ = 10 μm. A-H and A’-H’ were raised at 29°C and 25°C, respectively. Transgenes were expressed via *GMR-GAL4* driver; ATXN1 [82Q]/No modifier control used a neutral UAS line. (**I-L**) Motor performance quantification in flies of the indicated genotypes, measured as speed as a function of age. Control flies (blue lines) display better motor function across 25 days, while ATXN1 [82Q] expression in the nervous system (gray lines) causes progressive motor decline. Coexpression of KPNAs with ATXN1 [82Q] (red lines) mitigates this impairment. Dark lines represent regression splines; shaded areas indicate confidence intervals. Statistical significance between ATXN1 [82Q] alone (gray line) or together with a candidate gene overexpression (red line) is based on a non-linear mixed effect model ANOVA (α = 0.05) followed by Holme’s adjustment. (**I-L**) Flies were raised at 28°C. Transgenes were expressed via *nrv2-GAL4* driver. A neutral UAS line for ATXN1 [82Q]/No modifier control.

### Increased KPNA levels reduce overall nuclear ATXN1 [82Q] accumulation

Prior studies show that the interaction between KPNB and proteins with PrLDs that are mislocalized to the cytoplasm (like TDP-43, FUS, EWSR1, TAF15, hnRNPA1, and hnRNPA2) results in the defibrilization and nuclear relocalization of these proteins. ^10,20,25,26,36^. KPNB not only returns TDP-43 to the nucleus but can also suppress TDP-43-induced neurodegeneration ^26^. However, in the case of ATXN1 [82Q], nuclear localization of the protein is linked to its toxic effects, since mutating the NLS in polyQ-expanded ATXN1 eliminates its pathogenicity in mice ^25,31^. Thus, the neuroprotective effects of elevated KPNA levels are unlikely attributable to increased nuclear import of ATXN1 [82Q]. Other potential mechanisms of neuroprotection have been proposed for KPNs, including evidence supporting chaperoning and disaggregation functions ^20,25,37^. Our immunofluorescence assays revealed that ATXN1 [82Q] physically interacts with KPNAs in the *Drosophila* CNS. Therefore, we performed ATXN1 immunofluorescence assays in animals expressing ATXN1 [82Q] either alone or in combination with the KPNAs that ameliorated neurodegeneration. We analyzed both the level and nucleocytoplasmic localization of ATXN1. Surprisingly, we found that increased levels of either *Drosophila* KPNA6 (Kap-α1) (Fig. **3C**) or human KPNA2 (Fig. **3D**) resulted in decreased nuclear localization of ATXN1 [82Q] and increased accumulation of ATXN1 in the cytoplasm (Fig. **3C-D**). Quantification of confocal images confirmed that ATXN1 intensity was lower in the nuclei of neurons co-expressing ATXN1 [82Q] together with KPNAs (Fig. **3E-F**). These same neurons displayed increased cytoplasmic ATXN1 signal (Fig. **3G-H**). Given that increased KPNA levels ameliorated neurodegeneration, these results suggest that rather than increasing nuclear translocation, KPNA expression caused mutant ATXN1 [82Q] to concentrate in the cytoplasm, preventing its deleterious effects in the nucleus.

**FIG. 3.**
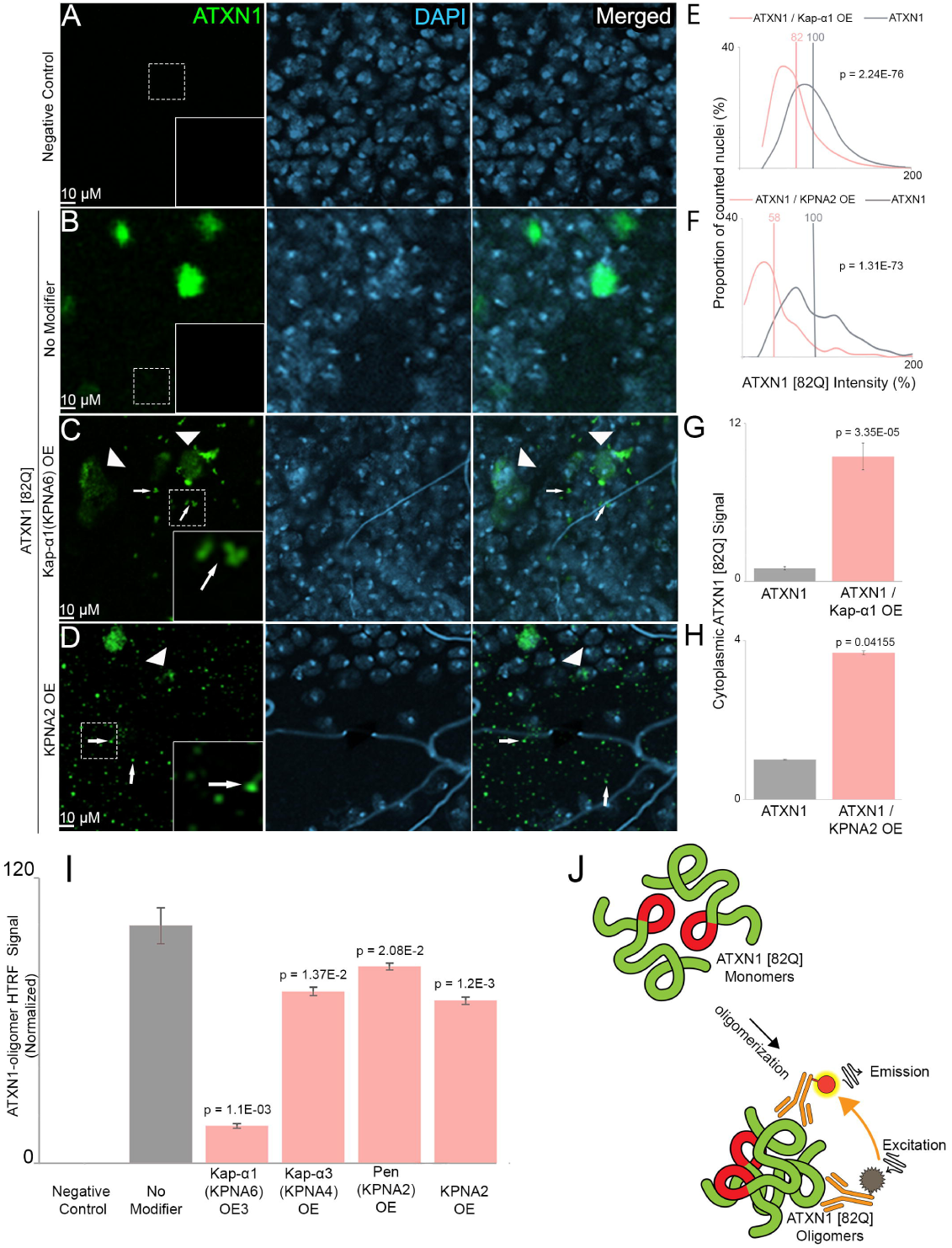
Increased KPNA levels reduce nuclear ATXN1 [82Q] accumulation and soluble oligomeric ATXN1 [82Q]. (**A-D**) Confocal images of *Drosophila* neurons in the adult VNCs stained with anti-ATXN1 [82Q] antibody (green), and DAPI marking the nucleus (blue). (**A**) No ATXN1 [82Q] (green) is detected in the neurons of negative control animals. (**B**) ATXN1 [82Q] staining (green) detected in discrete aggregates predominantly localized to the nucleus (blue in merged). (**C-D**) Notice the decreased nuclear ATXN1 [82Q] signal (arrowheads) and increased cytoplasmic ATXN1 [82Q] signal (small arrows) when ATXN1 [82Q] is expressed alongside different KPNAs. (**E-F**) Histograms depicting the proportion of nuclei as a function of nuclear ATXN1 [82Q] intensity show a leftward shift (lower nuclear ATXN1) in KPNA-expressing flies (pink) compared to ATXN1 [82Q] alone (gray). Average nuclear ATXN1 levels were significantly reduced (vertical lines). (**G-H**) Bar graphs show increased cytoplasmic ATXN1 [82Q] upon KPNA coexpression. Scale bars A-D = 10 μm. Flies raised at 28°C. Transgenes expressed via *nrv2-GAL4*; controls used a neutral UAS line. Data in E-H from 5 VNCs/genotype. Statistical significance: ANOVA followed by Student’s t-test. Error bars: SEM. **(I)** Homogeneous Time-Resolved Fluorescence (HTRF) detects soluble ATXN1 [82Q] oligomers in fly heads. No oligomers in negative controls (Lane 1); oligomers present in ATXN1 [82Q]-expressing flies (Lane 2). Coexpression with KPNAs (Lanes 3-6) significantly reduces ATXN1 [82Q] oligomers. Flies raised at 29°C. (**J**) Schematic of the HTRF assay: the same antibody is labeled with Europium Cryptate and D2, producing a signal only when ATXN1 [82Q] molecules interact. Statistical significance: ANOVA with Dunnett’s post-hoc test. Error bars: SEM.

### Increased KPNA levels reduce ATXN1 [82Q] soluble oligomers

Our previous results indicate that increased KPNA levels may chaperone ATXN1 [82Q] and limit its translocation to the nucleus. As in other neurodegenerative diseases, soluble ATXN1 [82Q] oligomers are likely the most pathogenic form of aggregates ^38^. To test whether increasing KPNA levels would influence mutant ATXN1 oligomerization, we developed a Homogeneous Time Resolved Fluorescence (HTRF) assay (Fig. **3I**). In this assay HTRF signal will only be produced when two ATXN1 [82Q] monomers form a dimer or oligomer. Since a mild detergent (TritonX) is used and the aqueous supernatant is tested, the assay will only be sensing the presence of soluble oligomers. We found that increased levels of the *Drosophila* homologs of KPNA2 (pen), KPNA4 (Kap-α3) and KPNA6 (Kap-α1) as well as human KPNA2, resulted in a decrease in ATXN1 [82Q] soluble oligomer HTRF signal detected in the *Drosophila* head extracts (Fig. **3I**). This result supports that KPNAs prevent ATXN1 [82Q] from adopting potentially pathogenic oligomeric conformations.

### KPNB1 inhibition elevates cytoplasmic KPNA, reduces nuclear mATXN1, and alleviates ATXN1 [82Q]-induced neurodegeneration

Given that increased KPNA levels ameliorate mutant ATXN1-induced neurodegeneration without enhancing its nuclear import, we posited that KPNA exerts a non-canonical function, chaperoning mATXN1 in the cytoplasm. In classical nuclear import, KPNA recognizes the substrate’s NLS and interacts with KPNB to facilitate transport into the nucleus ^39^. This led us to investigate if inhibiting KPNB would elevate cytoplasmic KPNA and reduce nuclear levels of mATXN1. We treated DAOY cells with INI-43, an inhibitor of KPNB1 ^40^, to assess how pharmacological inhibition of KPNB1 alters nucleocytoplasmic distribution of KPNAs. We found that INI-43 treatment resulted in increased KPNA2, KPNA5, and KPNA6 levels in the cytoplasm of DAOY cells as compared to DMSO control treatment (Fig. **4A-F**). Consistent with the observed increase in cytoplasmic KPNAs, we found that inhibition of KPNB1 with INI-43 resulted in increased levels of cytoplasmic ATXN1 [82Q] and decreased levels of nuclear ATXN1 [82Q] (Fig. **4G-J**). These findings indicate that KPNB1 inhibition increases cytoplasmic KPNA levels and promotes cytoplasmic retention of ATXN1 [82Q]. Taken together with our previous findings, these results suggest that the increased cytoplasmic KPNA due to KPNB1 inhibition is retaining and chaperoning mutant ATXN1 in the cytoplasm. We next examined whether KPNB1 inhibition would similarly modulate ATXN1 [82Q] localization and toxicity in our *Drosophila* model. We found that co-expression of ATXN1 [82Q] along with knockdown alleles of the *Drosophila* homolog of KPNB1, (Fs(2)Ket), ameliorated the compound eye and retinal photoreceptor degeneration characteristic of mATXN1 expression (Fig. **5A-D; 5A’-D’;** Supporting Information **S2**), and significantly attenuated the ATXN1 [82Q]-induced progressive neuronal dysfunction (Fig. **5E-G**). Lastly, immunofluorescence of the *Drosophila* VNC revealed that Fs(2)Ket (KPNB1) knockdown reduces nuclear levels of mutant ATXN1 (Fig. **5H-K**). These results suggest that KPNB1 knockdown mitigates ATXN1 [82Q]-induced neurodegeneration in *Drosophila* by reducing its nuclear accumulation.

**FIG. 4.**
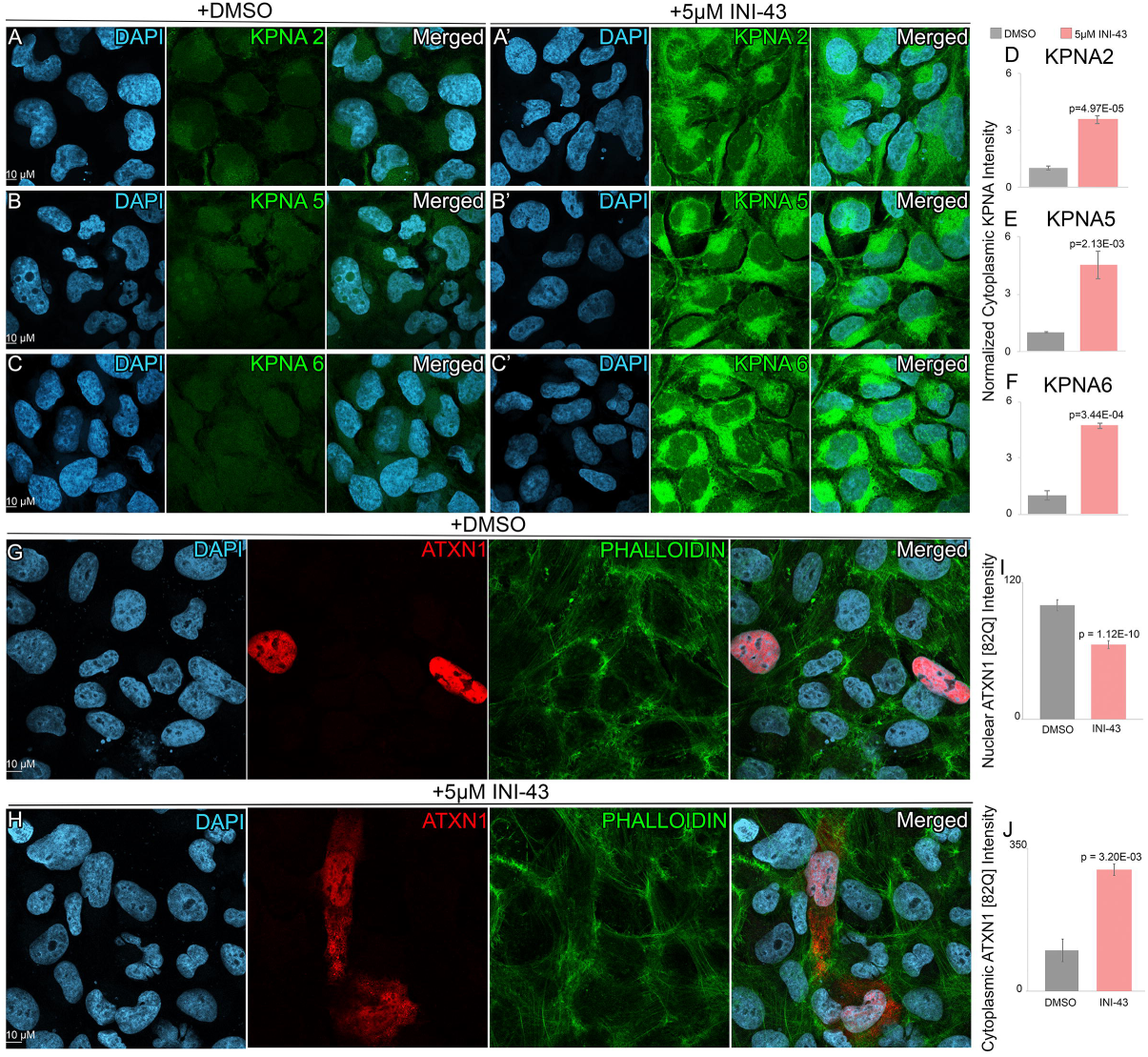
Pharmacological inhibition of KPNB1 increases cytoplasmic levels of KPNAs and alters nucleocytoplasmic distribution of mutant ATXN1. (**A-C’**) Confocal images of DAOY cells treated with DMSO (0.5%) and a KPNB1 inhibitor (INI-43 5µM) for 48 hours. Anti-KPNA2, Anti-KPNA5, and Anti-KPNA6 antibodies (green) and DAPI marking the nuclei (blue) indicate that compared to DMSO (0.5%) treatment (**A, B, C**), the inhibition of KPNB1 using 5µM INI-43 increased the cytoplasmic levels of KPNA2, KPNA5, and KPNA6 (**A’, B’, C’**). (**D-F**) Bar graphs depict a significant increase in cytoplasmic intensity of KPNA2, KPNA5, and KPNA6 (pink bars) when DAOY cells are treated with INI-43 as compared to DMSO control treatment (gray bars). (**G and H**) Confocal images of DAOY ATXN1 [82Q]-RFP cells stained with anti-ATXN1 [82Q] antibody (red) and DAPI marking the nuclei (blue); phalloidin (green) is used as a cytoplasmic counterstain. (**G**) Control treatment with DMSO (0.5%) for 24hrs shows ATXN1 [82Q] staining (red) is detected in discrete nuclear aggregates. **(H)** Treatment for 24 hours with 0.5uM INI-43 shows an increased cytoplasmic localization and decreased nuclear localization of ATXN1 [82Q] (red). **(I and J)** Bar graphs depicting quantification of average nuclear ATXN1 [82Q] intensity in DAOY ATXN1 [82Q]-RFP cells. Notice the significant decrease in nuclear ATXN1 [82Q] intensity concordant with an increase in cytoplasmic ATXN1 [82Q] intensity in the INI-43 treated cells (pink bars) as compared to control cells treated with DMSO (gray bars). P values were determined using ANOVA followed by Student’s t test (α = 0.05). Error bars are standard errors of the mean. Scale bars = 10μm.

**Fig. 5.**
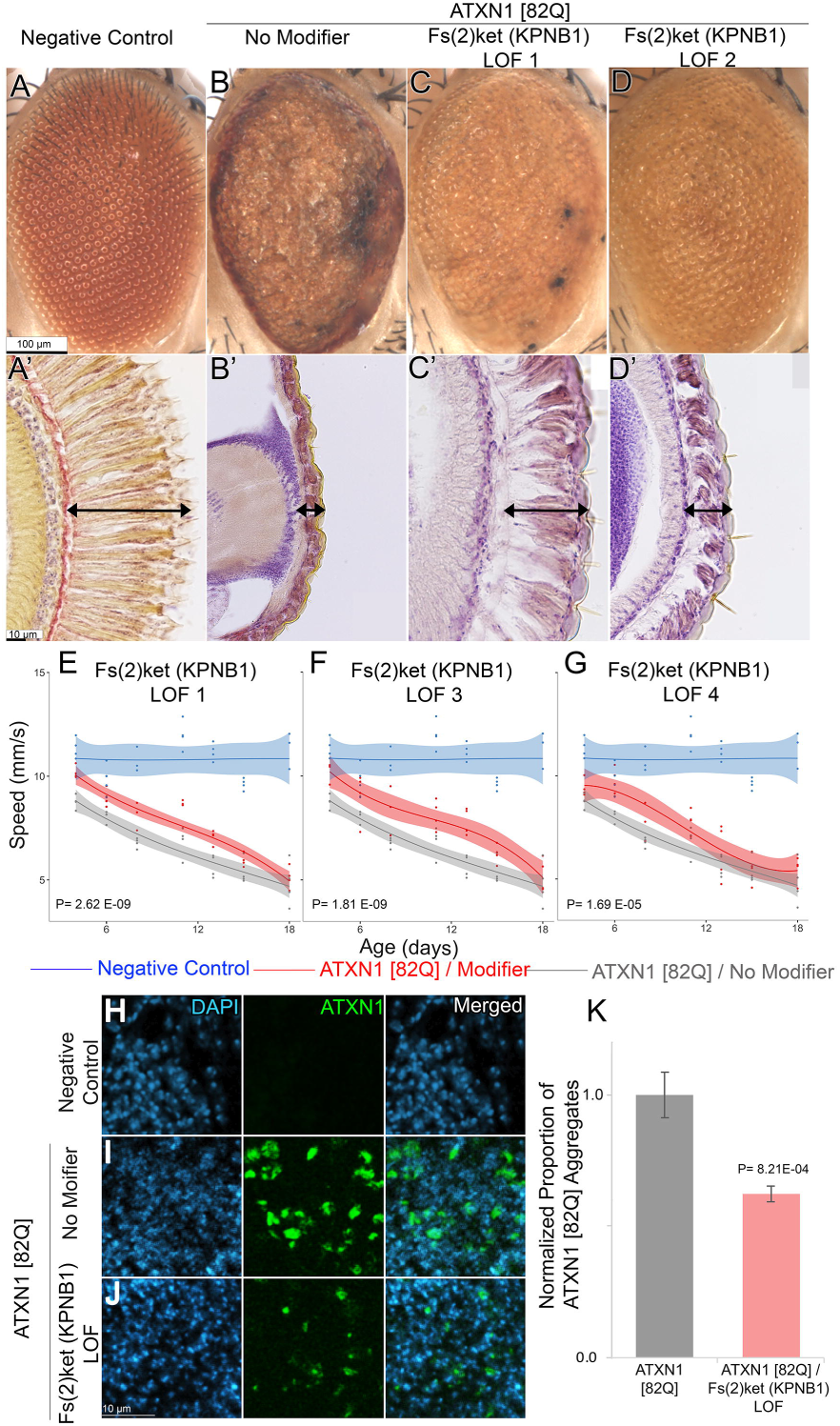
KPNB1 knockdown ameliorates ATXN1 [82Q]-induced neurodegeneration and reduces nuclear aggregation. (**A–D**) External eye images and (**A’–D’**) retinal paraffin sections of indicated genotypes. (**A, A’**) Wild-type controls show organized ommatidia and evenly spaced interommatidial bristles (A), and long, parallel photoreceptors without gaps (A’). Arrows indicate photoreceptor length. (**B, B’**) ATXN1 [82Q] expression causes disorganized ommatidia, missing bristles (B), and short, curved photoreceptors (B’). (**C–D, C’-D’**) Knockdown of *Drosophila* homolog of KPNB1 improves ommatidial structure (C–D) and photoreceptor length and alignment (C’–D’) compared to ATXN1 [82Q] alone. Driver was *GMR-GAL4* and a neutral UAS line was used for the ATXN1 [82Q]/No modifier control. (**E–G**) Quantification of motor performance (speed in mm/s) over time (days) in flies expressing ATXN1 [82Q] alone or with KPNB1 knockdown in the CNS. Age-matched trials show that wild-type controls (blue lines) perform significantly better compared to control flies expressing just ATXN1 [82Q] (gray lines) in the CNS. Co-expression of ATXN1 [82Q] in the CNS together with knockdown of *Drosophila* homologs of KPNB1 ameliorates ATXN1 [82Q]-induced motor impairments (red lines). Driver was *nrv2-GAL4* and a neutral UAS line was used for the ATXN1 [82Q]/No modifier control. (**H-J**) Confocal images of adult *Drosophila* VNC neurons stained for ATXN1 [82Q] (green) and nuclei (DAPI, blue). (**H**) No ATXN1 [82Q] detected in negative control. (**I**) ATXN1 [82Q] (green) forms nuclear aggregates. (**J**) Coexpression of mutant ATXN1 with knockdown of *Drosophila* homolog of KPNB1 reduces nuclear aggregates. (**K**) Bar graphs show decreased nuclear ATXN1 [82Q] upon coexpression of KPNB1 knockdown. Scale bars: A–D = 100 μm; A’–D’ & H-J = 10 μm. Flies were raised at: 29°C in A-D; 25°C in A’-D’, and 28°C E-K. Data in O from 5 VNCs/genotype. Statistical significance in E-G between ATXN1 [82Q] alone (gray line) and with candidate gene knockdown (red line) was assessed using a non-linear mixed-effect model ANOVA (α = 0.05). Statistical significance in K: ANOVA followed by Student’s t-test. Error bars: SEM.

## Discussion

The role played by NCT in neurodegeneration is an expanding area of research ^13,19^. On one hand, many disease driving proteins cause abnormalities in the distribution or function of NCT proteins or nuclear structures like the nuclear lamina, which are also observed in patients ^11,15,16,22^. On the other hand, nucleocytoplasmic mislocalization of disease-driving-proteins like TDP-43, Tau or Synuclein is observed in patient brains of AD, ALS, FTD or PD ^6,11,41^. In line with this, the abnormal nuclear or cytoplasmic localization of the disease driving proteins influences their toxicity ^25,31^.

Increasing KPNB levels has been previously shown to modulate the neurodegeneration caused by TDP-43 and ATXN3 ^20,26,42,43^. Here we show for the first time that increasing the levels of KPNAs ameliorates the neurodegeneration induced by mutant ATXN1. Interestingly, whereas the protective effect of increased KPNB levels is associated with its ability to relocate TDP-43 back to the nucleus ^26^, this is not the case in the KPNA-ATXN1 interaction. Rather, increased KPNA levels result in the retention of mutant ATXN1 in the cytoplasm and its decrease in the nucleus. Although previous studies have demonstrated that mutating the NLS sequence in mATXN1 is neuroprotective ^31,32^, our findings provide a proof of principle that genetic or pharmaceutical manipulations preventing the nuclear translocation of mATXN1—even when its NLS remains functional—should lead to neuroprotection in the context of SCA1. This points to an attractive therapeutic avenue.

It has been suggested that KPNs can act as chaperones with the ability to disaggregate disease driving proteins (like TDP-43) before transporting them to the nucleus ^10,20,25,26,36^. Our observations in the context of the ATXN1-KPNA interaction suggest that the protective chaperoning ability of KPNs can be uncoupled from the nuclear transport function. This is supported by the fact that we see decreased nuclear ATXN1 and increased cytoplasmic ATXN1 when the levels of KPNA are increased. Additionally, we observe a decrease in ATXN1 soluble oligomers when the levels of KPNAs are increased. Taken together, our results suggest that the binding of mATXN1 and KPNA results in non-toxic cytoplasmic retention and a decrease in toxic oligomers.

Our findings that KPNB1 inhibition increases cytoplasmic KPNA levels and reduces nuclear accumulation of mutant ATXN1 in DAOY cells suggest that elevated KPNA levels are neuroprotective in the context of mATXN1. This is further supported by the observed amelioration of ATXN1 [82Q]-induced neurodegeneration in *Drosophila* by KPNB1 knockdown. Inhibition of KPNB1 itself may prevent nuclear translocation of mutant ATXN1, thereby preventing its neurotoxicity. Our results also suggest an additional mechanism: inhibition of KPNB1 increases cytoplasmic KPNAs potentially chaperoning ATXN1 [82Q] and preventing formation of pathogenic oligomers. Though previous research has shown that KPNA may also translocate into the nucleus in a KPNB-independent manner, this nuclear import of KPNA is saturable and elevated levels of KPNA result in its cytoplasmic retention ^39^.

Several neurodegeneration-driving proteins including mATXN1 have the capacity to interfere with NCT ^4,11,22,30^. While restoration of impaired NCT can be a potential therapeutic strategy for other neurodegenerative diseases, our findings that increased KPNA levels do not increase nuclear transport of mATXN1, as well as the neuroprotective effect of KPNB1 knockdown, argue against this being the main mechanism of action here.

Though outside the scope of this work, our future research will assess if disrupting NCT by knocking down other key components of NCT like Ran, RanGAP, and nucleoporins will provide similar neuroprotective effects.

While our findings support a protective role for KPNA-mediated cytoplasmic retention of mutant ATXN1, the precise mechanism by which KPNAs reduce ATXN1 oligomerization remains unresolved. Our models— *Drosophila* and cultured human cells—provide strong *in vivo* and *in vitro* evidence, but further validation in mice and human neuronal systems is needed. Additionally, the specificity and *in vivo* applicability of KPNB1 inhibitors like INI-43 remain to be established.

Tau and α-Synuclein are normally cytoplasmic proteins but have been observed in the nuclei of tauopathy and PD neurons respectively ^11,41,44^. Furthermore, their nuclear translocation, mediated by TRIM28 for example, results in exacerbated toxicity ^45^. Additionally, Tau has been shown to interact with NUP98, which promotes its aggregation ^11^. Alterations in KPNA and KPNB levels, as well as their nucleocytoplasmic distributions, are characteristic of many diseases including AD and PD. ^20,46^. This suggests that the neuroprotective role of KPNAs may extend beyond our observations in the context of SCA1, having broader therapeutic implications in the context of neurodegenerative diseases.

## Materials and methods

*Drosophila strains*: Alleles expressing the different KPNA isoforms or knockdown of KPNB1 used were obtained from the Bloomington Drosophila Stock Center (BDSC). The *UAS-ATXN1 [82Q] F7* strain was used for all experiments ^47^. *GMR-GAL4* (Freeman et al., 1996) was used to drive expression of ATXN1 [82Q] in the retina while *nrv2-GAL4* (BDSC line 6800) was used to drive ATXN1 [82Q] in the nervous system. The BDSC stock number of the alleles used for our experiments are recorded in Supporting Information **S1**.

### Immunofluorescence

Ten-day old *Drosophila* CNS was dissected in cold PBS and fixed in 4% formaldehyde/PBS for 25 mins at room temperature. DAOY cells were fixed in 4% formaldehyde/PBS for 15 mins at room temperature. Tissue and cells were incubated in primary antibody overnight at 4^°^C. We used 11NQ anti-ATXN1 as previously described ^48^, rat anti-HA (Sigma, clone 3F10, 1:400), Rabbit anti-KPNA2 (Sigma, HPA041270, 1:50) Rabbit anti-KPNA5 (Sigma, HPA054037, 1:50), and Rabbit anti-KPNA6 (Sigma, HPA018863, 1:50). Following washes, Alexa Fluor 488 and Alexa Fluor 555 secondary antibodies (Thermo Fisher Scientific) were used and Dapi was added to stain the nuclei. Images were captured using a Leica SP8 confocal system.

In order to quantify the nucleocytoplasmic distribution of ATXN1, we used ImageJ to create a mask using the Dapi signal. This allowed for the separate quantification of nuclear and cytoplasmic ATXN1 signal. The ImageJ Analyze Particles plugin was used to quantify the individual signal of each nucleus in order to generate the distribution.

### External eye images and retinal sections

For external eye images, the fruit flies were raised at 29^°^C. Images were captured using the Leica MZ16 imaging system. For retinal sections, animals were fixed (85% ethanol, 3.7% Formaldehyde, 5% Acetic Acid) overnight at room temperature. The tissue was dehydrated, embedded in paraffin and sectioned into 10 μm slices. Following deparaffinization and rehydration, slides were stained using Harris hematoxylin and mounted using Permount medium.

### Drosophila motor performance assay

To assess motor performance of fruit flies as a function of age, we used ten age-matched females per replica per genotype as previously described ^49^. Flies were collected over a 24-hour period and transferred into a new vial containing 300 μl of media every day. Animals were kept at 28^°^C. Four replicates were used per genotype. Using an automated platform, the animals are taped to the bottom of a plastic vial and recorded for 7.5s as they climb back up on the walls of the vials. Videos are analyzed using custom software to assess the speed of each individual animal. Four trials per replicate are performed each day shown, and four replicates per genotype are used. Using the average performance of all 10 animals in each replicate and 4 replicates per genotype, a nonlinear random mixed effect model ANOVA was applied to the average using each four replicates to establish statistical significance across genotypes. P-values were adjusted for multiplicity using Holm’s procedure. Code for this analysis is available upon request. All graphing and statistical analyses were performed in R. A neutral UAS line was used to generate negative controls to establish the healthy baseline motor performance and disease controls for the disease baseline.

### Homogeneous Time Resolved Fluorescence

11NQ antibody was labeled (Cisbio-Revvity) with Europium Cryptate (donor fluorophore) or with D2 (acceptor fluorophore). 42 heads per genotype were homogenized in 60ul of extraction buffer (1% TritonX in PBS + Protease inhibitor) and incubated at 4^°^C overnight. Readings were performed in triplicates of 20ul containing a 1:50 dilution of the extracts and 2ul of each labeled antibody. HTRF was evaluated using a Tecan Infinite® 200 PRO following the setting recommended by Cisbio-Revvity. In addition to the ATXN1 oligomerization HTRF signal calculation, controls of the exact same genotype but not expressing ATXN1 were used as blanks to account for background fluorescence.

### Cell culture

DAOY human medulloblastoma stable cell lines expressing mRFP-tagged ATXN1 [82Q] were used as previously described ^34^. INI-43, a KPNB1 inhibitor ^40,50^ was used at a 5uM concentration for 24 hours and 48 hours. Cells were fixed and stained as described above.

**Supplementary Materials 2. Knockdown of *Drosophila* homolog of KPNB1 ameliorates ATXN1 [82Q]-induced retinal degeneration**. (**A**) Wild-type control shows long and parallel photoreceptors without gaps. (**B**) ATXN1 [82Q] expression leads to short, curved photoreceptors (**C-D**) Expression of ATXN1 [82Q] together with knockdown of *Drosophila* homologs of KPNB1 ameliorates ATXN1 [82Q]-induced retinal degeneration. Arrows indicate photoreceptor length. Scale bars: A–D = 10 μm.

## Notes

### Competing Interest Statement

The authors have declared no competing interest.

### Summary of Updates

Figures 1-5 were out of order

