## Supplementary figures and images for "Modulation of Karyopherin Levels Attenuates Mutant Ataxin-1-Induced Neurodegeneration"

### Supplemental Figure 1

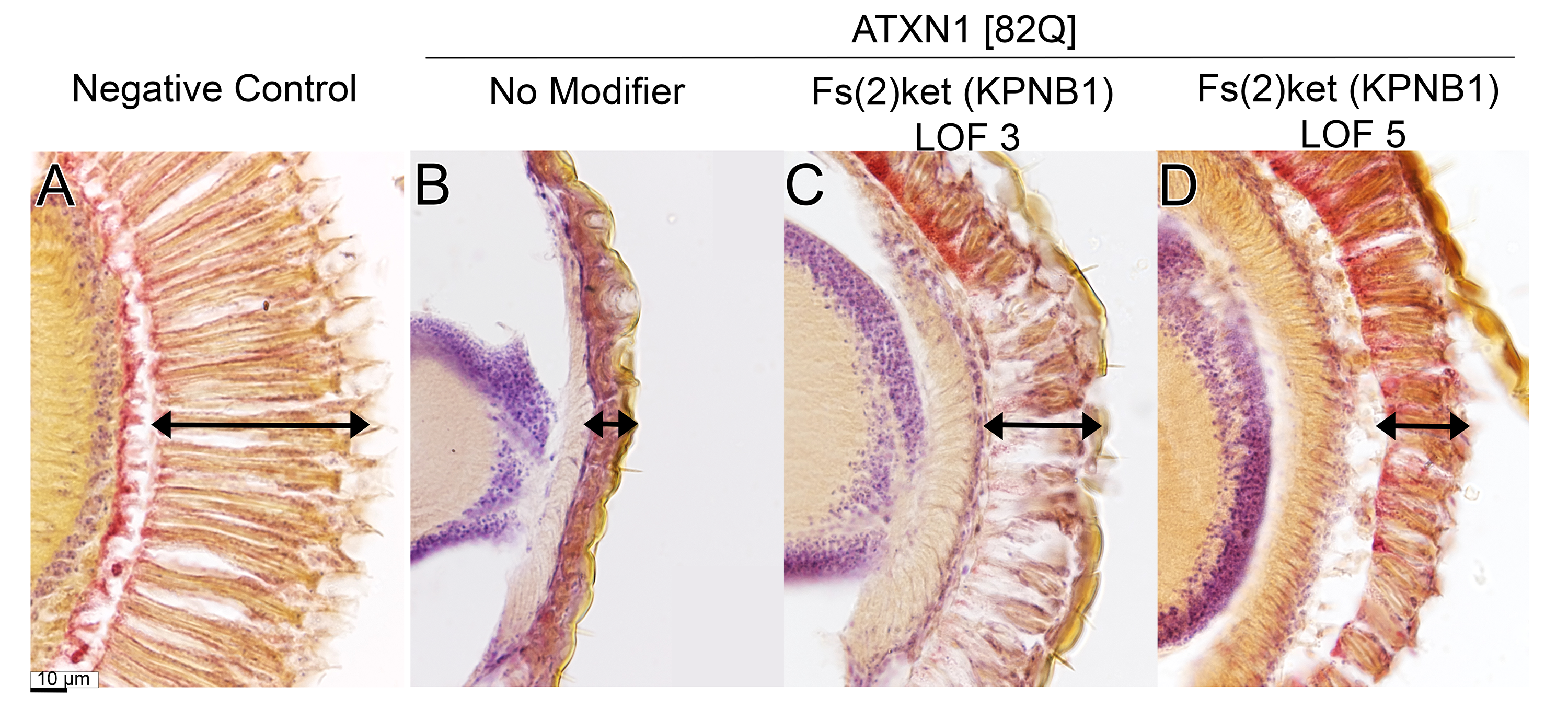
