## Supplemental Table 1 for "Modulation of Karyopherin Levels Attenuates Mutant Ataxin-1-Induced Neurodegeneration"

Supplemental Table 1- Allele identification for each figure.

| **Figure** | **#** | **Allele Identification** | ***Drosophila* Gene** | **Human Gene Correspondence** |
| --- | --- | --- | --- | --- |
| **Figure 1** | **A** | w; nrv2-GAL4 | N/A | N/A |
|  | **B** | w/UAS-ATXN1[82Q](F7); nrv2-GAL4 | N/A | N/A |
|  | **C** | BDSC-80688 | Kap-α1 | Homolog of Human KPNA6 |
|  | **D** | BDSC-91173 | N/A | Homolog of Human KPNA5 |
|  | **E** | BDSC-91180 | N/A | Homolog of Human KPNA2 |
| **Figure 2** | **A, A’** | w; GMR-GAL4 | N/A | N/A |
|  | **B, B’** | w/UAS-ATXN1[82Q](F7); GMR-GAL4 | N/A | N/A |
|  | **C, C’** | BDSC-27145 | Kap-α1 | Homolog of Human KPNA6 |
|  | **D, D’** | BDSC-25399 | Kap-α1 | Homolog of Human KPNA6 |
|  | **E, E’, I** | BDSC-15422 | Kap-α3 | Homolog of Human KPNA4 |
|  | **F, F’, J** | BDSC-20036 | Pen | Homolog of Human KPNA2 |
|  | **G, G’, K** | BDSC-91173 | N/A | KPNA5 (Human Gene) |
|  | **H, H’, L** | BDSC-91180 | N/A | KPNA2 (Human Gene) |
| **Figure 3** | **A** | w; nrv2-GAL4 | N/A | N/A |
|  | **B** | w/UAS-ATXN1[82Q](F7); nrv2-GAL4 | N/A | N/A |
|  | **C, E, G** | BDSC-27145 | Kap-α1 | Homolog of Human KPNA6 |
|  | **D, F, H** | BDSC-91180 | N/A | KPNA2 (Human Gene) |
|  | **I**- Kap-α1(KPNA6) OE3 | BDSC-80688 | Kap-α1 | Homolog of Human KPNA6 |
|  | **I**- Kap-α3(KPNA4) OE | BDSC-15422 | Kap-α3 | Homolog of Human KPNA4 |
|  | **I**- Pen(KPNA2) OE | BDSC-20036 | Pen | Homolog of Human KPNA2 |
|  | **I**- KPNA2 OE | BDSC-91180 | N/A | KPNA2 (Human Homolog) |
| **Figure 5** | **A, H** | w; GMR-GAL4 |  |  |
|  | **B, I** | w/UAS-ATXN1[82Q](F7); GMR-GAL4 |  |  |
|  | **C, C’, E** | BDSC-85029 | Fs(2)ket | Homolog of Human KPNB1 |
|  | **D, D’** | BDSC-35515 | Fs(2)ket | Homolog of Human KPNB1 |
|  | **F** | BDSC-44576 | Fs(2)ket | Homolog of Human KPNB1 |
|  | **G, J, K** | BDSC-4994 | Fs(2)ket | Homolog of Human KPNB1 |
| **Supplemental Figure 1** | **A** | w; GMR-GAL4 |  |  |
|  | **B** | w/UAS-ATXN1[82Q](F7); GMR-GAL4 |  |  |
|  | **C** | BDSC-44576 | Fs(2)ket | Homolog of Human KPNB1 |
|  | **D** | BDSC-27567 | Fs(2)ket | Homolog of Human KPNB1 |

OE=overexpression, LOF=loss of function
